# A neuron-optimized CRISPR/dCas9 activation system for robust and specific gene regulation

**DOI:** 10.1101/371500

**Authors:** Katherine E. Savell, Svitlana V. Bach, Morgan E. Zipperly, Jasmin S. Revanna, Nicholas A. Goska, Jennifer J. Tuscher, Corey G. Duke, Faraz A. Sultan, Julia N. Burke, Derek Williams, Lara Ianov, Jeremy J. Day

## Abstract

Recent developments in CRISPR-based gene editing have provided new avenues to interrogate gene function. However, application of these tools in the central nervous system has been delayed due to difficulties in transgene expression in post-mitotic neurons. Here, we present a highly efficient, neuron-optimized dual lentiviral CRISPR-based transcriptional activation (CRISPRa) system to drive gene expression in primary neuronal cultures and the adult brain of rodent model systems. We demonstrate robust, modular, and tunable induction of endogenous target genes as well as multiplexed gene regulation necessary for investigation of complex transcriptional programs. CRISPRa targeting unique promoters in the complex multi-transcript gene *Brain-derived neurotrophic factor* (*Bdnf*) revealed both transcript- and genome-level selectivity of this approach, in addition to highlighting downstream transcriptional and physiological consequences of *Bdnf* regulation. Finally, we illustrate that CRISPRa is highly efficient *in vivo*, resulting in increased protein levels of a target gene in diverse brain structures. Taken together, these results demonstrate that CRISPRa is an efficient and selective method to study gene expression programs in brain health and disease.

---

GENE EXPRESSION PATTERNS define neuronal phenotypes and are dynamic regulators of neuronal function in the developing and adult brain^1–3^. During development, differential expression of transcription factors induces gene programs responsible for neuronal fate specification and maturation^4^. In the adult brain, specific gene programs are altered by neuronal activity and behavioral experience, and these changes are critical for adaptive behavior^5–7^. Dysregulation of both developmental and adult brain gene programs is implicated in numerous neuropsychiatric diseases, such as addiction^8^, depression^9^, schizophrenia^10^, and Alzheimer’s disease^11^. Interrogating the role of gene expression programs in neuronal function has traditionally relied on the use of overexpression vectors^12^, transgenic animal models^13^, and knockdown approaches such as RNA interference^14^. While valuable, these techniques do not manipulate the endogenous gene locus, often require costly and time-consuming animal models, and are generally limited to one gene target at a time. Thus, while next-generation sequencing has allowed unprecedented characterization of gene expression changes in response to experience or disease, efficient multiplexed transcriptional modulation to recapitulate these expression patterns has proven elusive.

Recent advances in CRISPR/Cas9 genome editing have enabled unparalleled control of genetic sequences^15–17^, transcriptional states^18,19^, and epigenetic modifications^20^. This system has been harnessed for gene-specific transcriptional regulation by anchoring transcriptional effectors to a catalytically dead Cas9 (dCas9) enzyme, targeted to a select genomic locus with the help of a single guide RNA (sgRNA). However, these advances have not been readily adapted in the central nervous system (CNS) due to limitations in transgene expression in post-mitotic neurons^20^. For example, reports using CRISPR-based technologies in neurons required the use of cumbersome techniques such as *in utero* electroporation^16^, direct Cas9 protein infusion^21^, or biolistic transfection^16^. More widespread techniques such as virus-mediated neuronal transduction have been sparsely reported for gene knockdown^22^ or activation^23,24^, but the selectivity and function of these tools have not been systematically tested in neuronal systems.

Here, we present a modular, neuron-optimized CRISPR/dCas9 activation (CRISPRa) system to achieve robust upregulation of targeted genes in neurons. We show that a neuron-specific promoter is more efficient at driving the expression of CRISPR components in neurons over general ubiquitous promoters. Fusion of a robust transcriptional activator to dCas9 enabled effective gene upregulation despite gene class and size in primary rat cortical, hippocampal, and striatal neuron cultures. Co-transduction of multiple sgRNAs enabled synergistic upregulation of single genes as well as coordinated induction of multiple genes. CRISPRa targeting individual transcript promoters in *Brain-derived neurotrophic factor* (*Bdnf*) – a complex gene involved in synaptic plasticity, learning and memory^25^– revealed highly specific *Bdnf* transcript control without impact at non-targeted variants, and demonstrated the efficacy of this approach for studying downstream transcriptional programs and physiological functions. Finally, we validated these tools for *in vivo* applications in the prefrontal cortex, hippocampus, and nucleus accumbens of the adult rat brain. Our results indicate that this neuron-optimized CRISPRa system enables specific and large-scale control of gene expression profiles within the CNS to elucidate the role of gene expression in neuronal function, behavior, and neuropsychiatric disorders.

## RESULTS

### Optimization of CRISPRa for neuronal systems

As highlighted by previous studies, dCas9 fusion systems containing the transcriptional activator VPR (comprised of VP64 (a concatemer of the herpes simplex viral protein VP16), p65 (a subunit of the transcription factor NF-κB), and Rta (a gammaherpesvirus transactivator)), drive expression of target genes to a much higher degree as compared to single transactivators such as VP64 or p65 alone^26–28^. To achieve high construct efficiency while balancing size constraints due to the large size of the dCas9-VPR construct (>5.5 kbp), we assembled dual lentivirus-compatible plasmid constructs (**Figure 1a**) for separate expression of dCas9-VPR and sgRNA scaffolds. The sgRNA construct co-expresses mCherry and allows for convenient verification of its expression with live cell imaging, while dCas9-VPR contains a FLAG-tag for construct expression validation through immunocytochemistry (ICC). For dCas9-VPR cassette expression, we cloned various promoters previously shown to drive transgene expression in neurons^29^, including the ubiquitous promoters EF1α (human elongation factor 1 alpha), PGK (human phosphoglycerate kinase), and CAG (a strong synthetic hybrid promoter), as well as the neuron-specific promoter SYN (human synapsin 1 promoter). Construct functionality was validated in HEK293T cells targeting the human *FOS* gene (**Figure 1b**). For all CRISPRa manipulations, a sgRNA targeting the bacterial *LacZ* gene paired with dCas9-VPR was used as a non-targeting control. dCas9-VPR expressed from all tested promoters successfully drove *FOS* mRNA 40 hours after transfection as measured by RT-qPCR. Before validating these constructs in rat primary neurons, we further validated rat-specific sgRNAs in C6 cells (a dividing rat glioma cell line) using nucleofection of dCas9-VPR and sgRNA plasmids targeting either *LacZ* or the rat *Fos* gene (**Figure 1c**). Similar to HEK293T cells, dCas9-VPR expressed from all promoters was capable of inducing *Fos* mRNA. Finally, for robust expression in transfection-resistant post-mitotic neurons, we generated lentiviruses expressing sgRNA and dCas9-VPR constructs driven by various promoters. Lentiviral packaging with all dCas9-VPR plasmids generated high-titer lentiviruses (minimum 8.29 × 10^9^ GC/ml) with the exception of CAG-dCas9-VPR, which was excluded from subsequent experiments. Neuronal cultures prepared from embryonic rat cortex were transduced with either EF1α, PGK, or SYN-driven dCas9-VPR lentiviruses alongside sgRNAs targeted to either the bacterial *LacZ* or the rat *Fos* gene on days *in vitro* 4 (DIV 4), and RNA was harvested on DIV 11. Surprisingly, despite transducing with the same multiplicity of infection, only the SYN-dCas9-VPR lentivirus resulted in robust induction of *Fos* mRNA (**Figure 1d**). Taken together, our RT-qPCR results across cell lines and primary neurons indicate that while dCas9-VPR can be driven by multiple promoters in other cell types, only the SYN promoter drives sufficient transgene expression to produce a functional effect in primary neuronal cultures.

**Figure 1.**
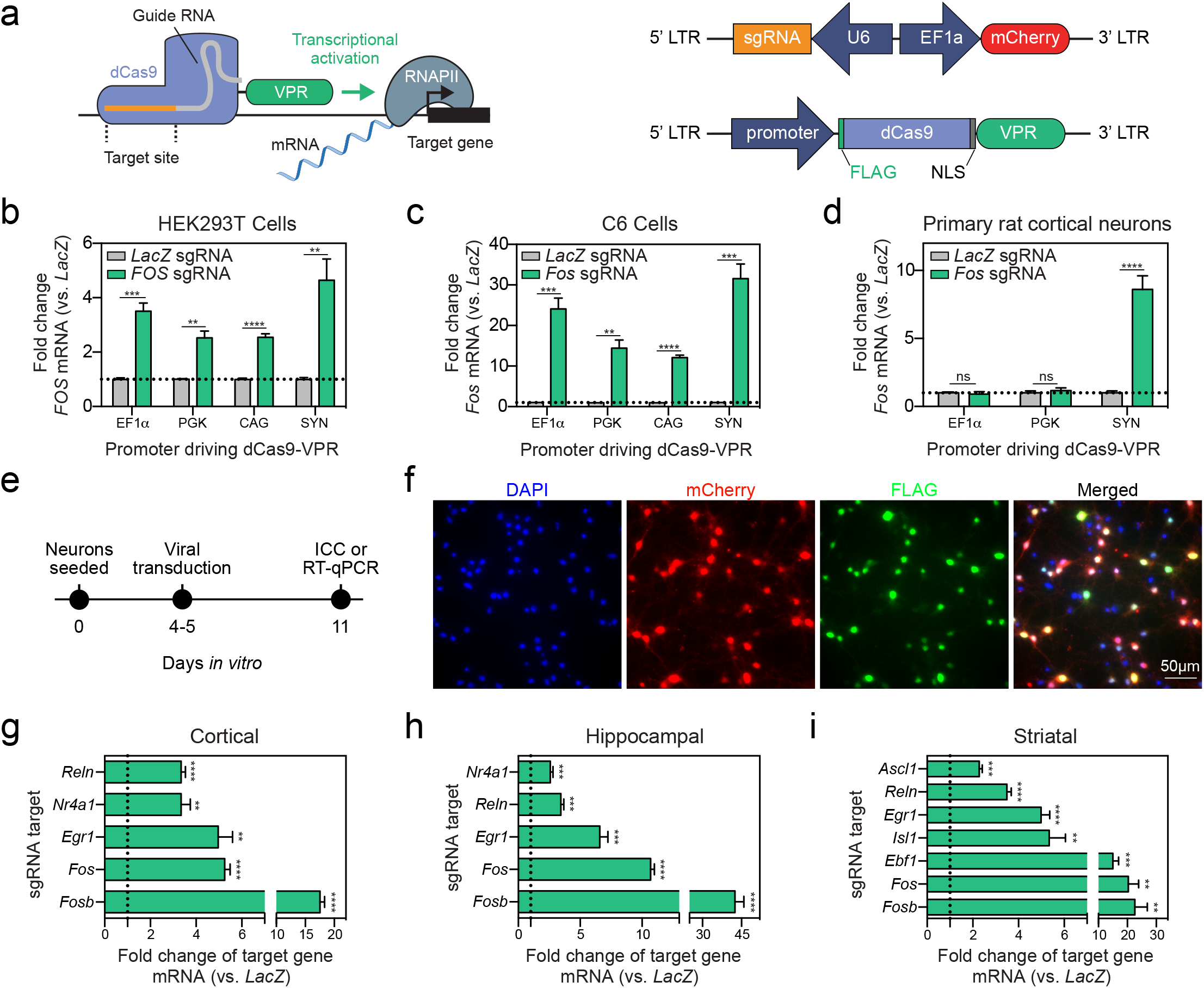
CRISPRa gene induction in HEK293T cells, C6 cells, and primary rat neurons under ubiquitous and neuron-selective promoters. (**a**) Illustration of the CRISPRa dual vector approach expressing either the single guide RNA (sgRNA) or the dCas9-VPR construct driven by EF1α, PGK, CAG, or SYN promoters. (**b**) dCas9-VPR co-transfected with sgRNAs targeted to the human *FOS* gene results in induction of *FOS* mRNA in HEK293T cells regardless of the promoter driving dCas9-VPR (*n* = 6, unpaired *t*-test; EF1α *t_5.308_* = 8.034, *P* = 0.0004; PGK *t_5.138_* = 5.943, *P* = 0.0018; CAG *t_6.097_* = 11.15, *P* < 0.0001; SYN *t_5.064_* = 4.67, *P* = 0.0053). (**c**) dCas9-VPR co-nucleofected with sgRNAs targeting the rat *Fos* gene induces *Fos* mRNA in a C6 glioblastoma cell line. (*n* = 6, unpaired *t*-test; EF1α *t_5.006_* = 8.699, *P* = 0.0003; PGK *t_5.067_* = 6.640, *P* = 0.0011; CAG *t_5.148_* = 18.32, *P* < 0.0001; SYN *t_5.000_* = 8.631, *P* = 0.0003). (**d**) Lentiviral transduction of primary rat cortical neurons reveals that only dCas9-VPR driven by the SYN promoter results in induction of *Fos* mRNA (*n* = 6, unpaired *t*-test; EF1α *t_6.912_* = 0.492, *P* = 0.6378; PGK *t_9.491_* = 0.710, *P* = 0.4950; SYN *t_5.234_* = 7.593, *P* = 0.0005). (**e**) Experimental timeline for *in vitro* CRISPRa in neurons. Primary rat neuronal cultures are generated and transduced with dual sgRNA/dCas9-VPR lentiviruses at days *in vitro* 4-5 (DIV 4-5). On DIV 11, neurons underwent either immunocytochemistry (ICC) to validate viral expression or RNA extraction followed by RT-qPCR to examine gene expression. (**f**), ICC reveals high co-transduction efficiency of guide RNA (co-expressing mCherry, signal not amplified) and dCas9-VPR (FLAG-tagged) lentiviruses in primary neuronal cultures. Cell nuclei are stained with 4,6-diamidino-2-phenylindole (DAPI). Scale bar, 50 μm. (**g-i**) dCas9-VPR increases gene expression for a panel of genes in cortical, hippocampal, or striatal cultures. Data are expressed as fold change of the target gene’s expression relative to dCas9-VPR targeted to a non-targeting control (bacterial *LacZ* gene). (*n* = 4-6, unpaired *t*-test; Cortical: *Reln t_5.438_* = 12.590, *P* < 0.0001; *Nr4a1 t_3.250_* = 5.692, *P* = 0.0086; *Egr1 t_5.084_* = 6.233, *P* = 0.0015; *Fos t_5.571_* = 16.770, *P* < 0.0001; *Fosb t_5.167_* = 19.570, *P* < 0.0001; Hippocampal: *Nr4a1 t_5.760_* = 7.140, *P* = 0.0005; *Reln t_6.102_* = 7.236, *P* = 0.0003; *Egr1 t_5.091_* = 8.565, *P* = 0.0003; *Fos t_6.668_* = 27.410, *P* < 0.0001; *Fosb t_5.021_* = 12.210, *P* < 0.0001; Striatal: *Ascl1 t_5.111_* = 9.383, *P* = 0.0002; *Reln t_5.667_* = 12.790, *P* < 0.0001; *Egr1 t_5.760_* = 10.320, *P* < 0.0001; *Isl1 t_5.047_* = 6.074, *P* = 0.0017; *Ebf1 t_5.012_*= 7.007, *P* = 0.0009; *Fos t_5.026_* = 5.349, *P* 0.003; *Fosb t_4.015_* = 5.057, *P* = 0.0071). dCas9-VPR with a sgRNA targeted to the bacterial *LacZ* gene is used as a non-targeting control in panels (**b-d**) and (**g-i**). All data are expressed as mean ± s.e.m. Individual comparisons, ^**^*P* < 0.01, ^***^*P* < 0.001 and ^****^*P* < 0.0001.

Different regions in the brain have diverse neuronal subtypes, so we next sought to validate whether the SYN-driven CRISPRa system could be utilized in neuronal cultures with differing neuronal composition. Primary cultures from embryonic cortex, hippocampus, or striatum were generated and transduced with the dual lentivirus CRISPRa system. On DIV 11, cultures were used for either ICC or RNA extraction to examine gene expression with RT-qPCR (**Figure 1e**). ICC revealed high co-localization of the sgRNA (co-expressing mCherry, signal not amplified) and the dCas9-VPR construct (FLAG-tagged) in cortical neurons (**Figure 1f**). To assess the efficacy of the CRISPRa system at multiple gene targets, we designed one to three sgRNAs per gene targeting promoter regions within 1.5 kbp to 100 bp upstream of the transcriptional start site (TSS) of a given target gene. We targeted an array of genes important to neuronal development, plasticity, and learning and memory, including immediate early genes (*Egr1*, *Fos*, *Fosb*, and *Nr4a1*), neuron-defining transcription factors (*Ascl1*, *Isl1*, *Ebf1*), and an extracellular matrix protein (*Reln*)^3–5^. These genes varied in length from 1.8 kbp (*Ascl1*) to 426.1 kbp (*Reln*). For each targeted gene, we found significant induction of gene expression compared to the *LacZ* non-targeting control (**Figure 1g-i**). Successful induction of a variety of targets, despite gene function or length, in multiple neuronal subpopulations suggests that this CRISPRa system can be used to drive gene expression at a large number of genes within the mammalian CNS, regardless of neuronal cell type.

### CRISPRa multiplexing enables synergistic and coordinated gene regulation

CRISPRa-mediated upregulation produced a range of magnitudes in induction between target genes. Therefore, to test whether targeting multiple copies of dCas9-VPR to a single gene boosted observed mRNA induction, we pooled between one and three sgRNA lentiviruses for each selected gene target (**Figure 2a**). We focused on the immediate early genes *Fos* (3 pooled sgRNAs) and *Fosb* (2 pooled sgRNAs), as they produced the most robust changes in gene expression in all neuronal subpopulations. For both *Fos* and *Fosb*, combining sgRNAs synergistically induced gene expression over an individual sgRNA (**Figure 2b**), suggesting that target gene induction can be titrated with CRISPRa to produce the desired level of gene induction.

**Figure 2.**
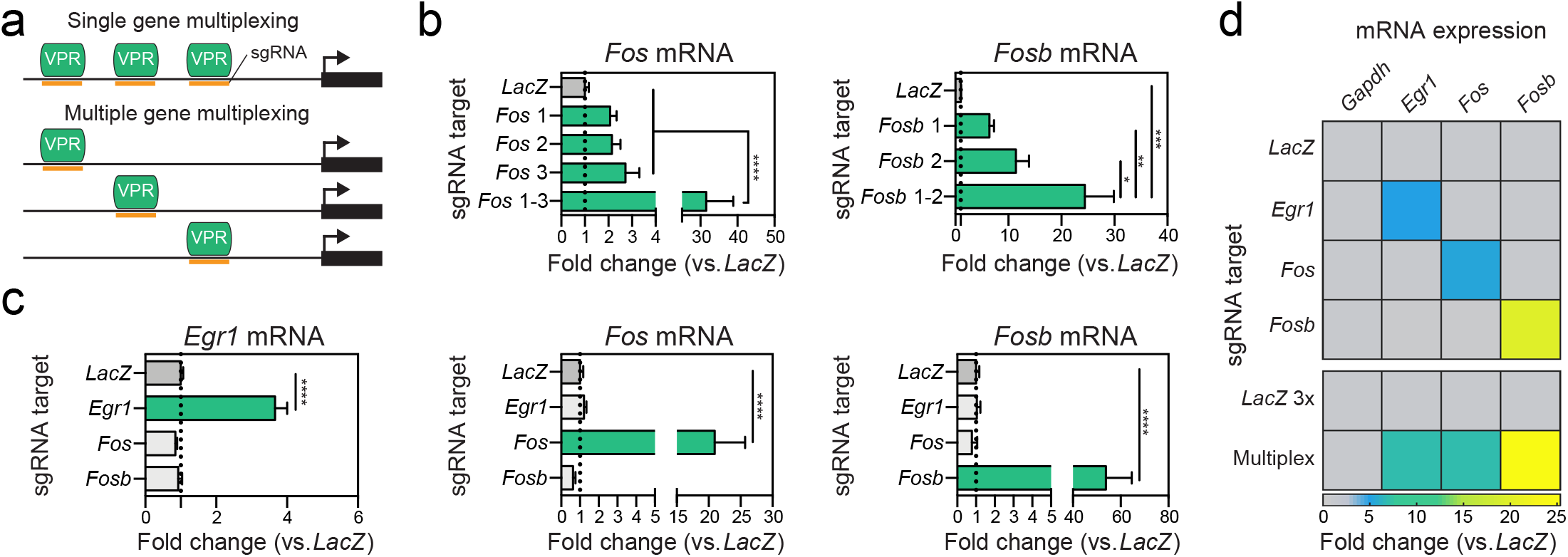
CRISPRa sgRNA multiplexing for synergistic or coordinated control of gene expression. (**a**) Illustration of pooled sgRNA multiplexing for dCas9-VPR targeting to multiple locations at a single gene (top) or simultaneous regulation of several genes (bottom). (**b**) Single gene multiplexing at *Fos* (left) and *Fosb* (right) reveals that while individual sgRNAs are sufficient to drive gene expression, sgRNA pooling results in synergistic induction of gene expression in cultured neurons (*n* = 5-6, one-way ANOVA, *Fos* F_(4,25)_ = 16.17, *P* < 0.0001; *Fosb* F_(3,19)_ = 10.23, *P* = 0.0003; Tukey’s *post hoc* test for individual comparisons). (**c**) CRISPRa with sgRNAs targeting *Egr1*, *Fos*, or *Fosb* individually results in specific and robust increases in gene expression without off-target effects. (*n* = 5-6, one-way ANOVA, *Egr1* F_(3,16)_ = 56.53, *P* < 0.0001; *Fos* F_(3,16)_ = 17.55, *P* < 0.0001; *Fosb* F_(3,15)_ = 32.06, *P* < 0.0001; Dunnett’s *post hoc* test for individual comparisons). (**d**) Pooled gRNAs result in coordinated increases in gene expression at *Egr1*, *Fos*, and *Fosb* (*n* = 6 per group). All data are expressed as mean ± s.e.m. Individual comparisons, ^*^*P* < 0.05, ^**^*P* < 0.01 ^***^*P* < 0.001 and ^****^*P* < 0.0001.

Next, we sought to investigate whether the CRISPRa system could be used to drive simultaneous expression of multiple genes, providing a method to study more coordinated changes in gene expression (**Figure 2a**). We focused on three immediate early genes (*Fos*, *Fosb*, and *Egr1*), all of which are rapidly induced after neuronal activity and have well-established roles in neuronal function and behavior^5^. First, we individually recruited dCas9-VPR to each gene’s promoter region in striatal cultures, which resulted in robust increases of gene expression without altering the baseline of the other genes (**Figure 2c**). Next, we combined the sgRNA lentiviruses for all three gene targets, which resulted in simultaneous induction of gene expression of all three genes (**Figure 2d**). While we have not tested the limit of how many genes can be simultaneously induced with this system, these results demonstrate that our CRISPRa system can be used to study complex gene expression programs that normally occur in response to neuronal activation.

Previous work has introduced a CRISPR interference (CRISPRi) system in neurons, in which the dCas9 is fused to a transcriptional repressor, KRAB^22^. We tested whether the same sgRNAs used in our CRISPRa system could also be used to repress the same gene target with CRISPRi (**Supplementary Figure 1a**). As previously described, sgRNAs that are close to the TSS are most effective for transcriptional repression, and we found that for *Egr1* and *Fosb*, KRAB-dCas9 targeting resulted in a blunting of gene expression (**Supplementary Figure 1b**). For *Fos*, which has sgRNAs designed at larger distances from the TSS, KRAB-dCas9 was not effective at reducing gene expression. Interestingly, we found that downregulating *Egr1* also affected baseline *Fosb* levels, suggesting that *Egr1* is necessary for *Fosb* expression. Taken together, it is possible that sgRNAs can be utilized for both the CRISPRa or CRISPRi systems to bidirectionally regulate gene expression.

### Selective upregulation of distinct transcript variants with CRISPRa

To examine the specificity of CRISPRa in neurons, we tested whether it is possible to drive transcription of a single transcript variant of a gene. We chose *Brain-derived neurotrophic factor* (*Bdnf*) as our target gene due to its complex transcriptional regulation and central role in diverse processes such as neuronal differentiation and survival, dendritic growth and synaptic development, long-term potentiation (LTP), and memory formation^21,27,28^. The *Bdnf* gene consists of nine 5’ non-coding exons (*I-IXa*) and one 3’ coding exon (*IX*) (**Figure 3a**)^30^. Each non-coding exon has its own unique upstream promoter region where transcription of each variant is initiated. Differential promoter usage gives rise to diverse transcripts that incorporate at least one non-coding 5’ exon in combination with the 3’ coding exon, all of which code for the same mature Bdnf protein^30^. Due to this complexity, attempts to characterize distinct functional roles of individual *Bdnf* mRNAs in neurons have produced conflicting results^31,32^, and currently available tools either lack the ability to selectively upregulate single *Bdnf* transcript variants or require cumbersome molecular cloning protocols to generate gene-specific targeting constructs.

**Figure 3.**
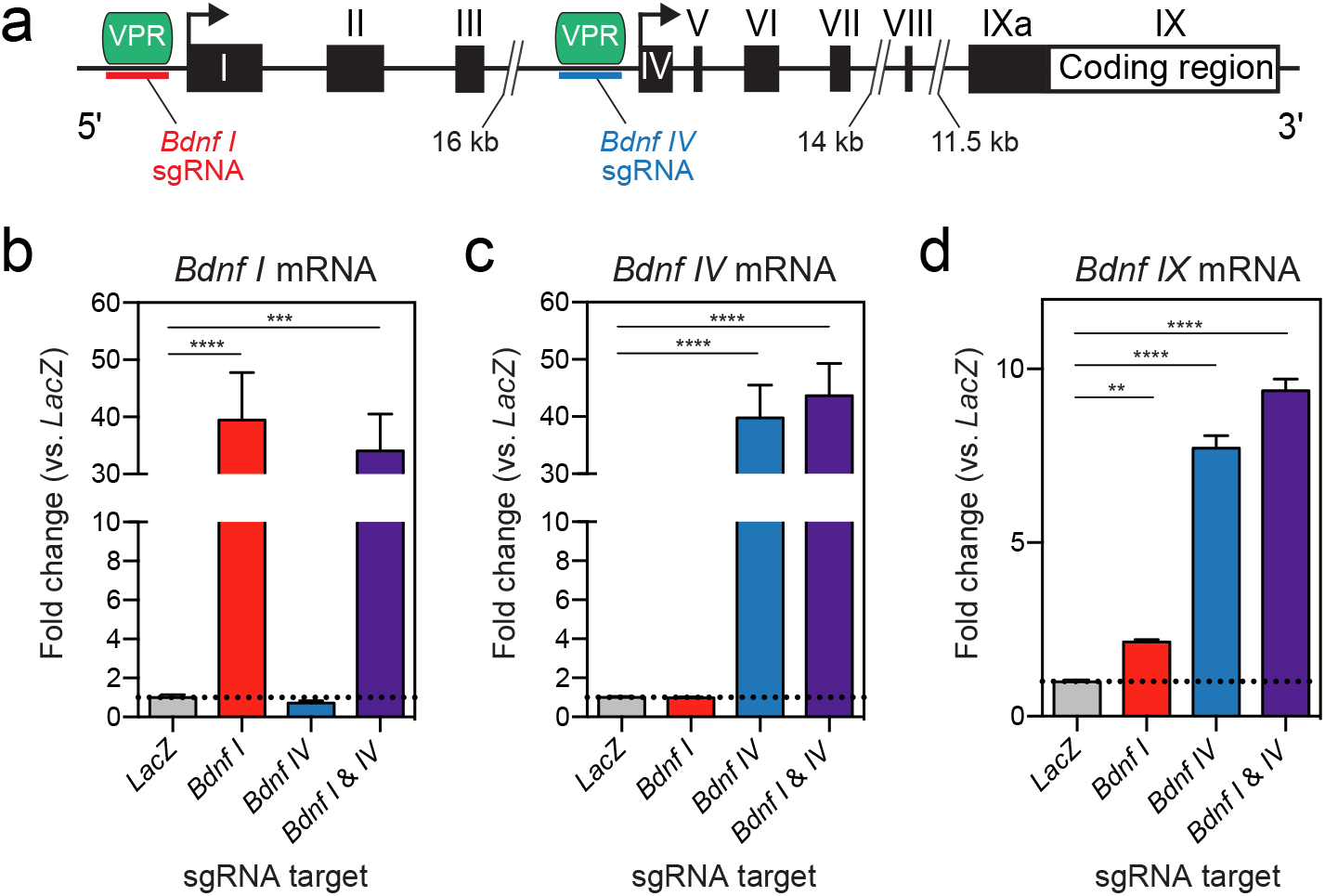
CRISPRa induction of *Bdnf* transcript variants *I* and *IV* in primary rat hippocampal neurons. (**a**) *Bdnf* gene structure illustrating non-coding exons (*I-IXa*) and a common coding exon (*IX*). sgRNAs were designed upstream of exons *I* and *IV*, as indicated by the red and blue lines. (**b-d**) Expression of *Bdnf I*, *IV* and *IX* transcript variants after targeting dCas9-VPR to exons *I* and/or *IV* using sgRNAs, measured with RT-qPCR. (**b**) *Bdnf I* transcript is specifically upregulated with *Bdnf I* sgRNA but not with *Bdnf IV* sgRNA (*n* = 8, one-way ANOVA, F_(3, 28)_ = 15.65, *P* < 0.0001). (**c**) *Bdnf IV* transcript is specifically upregulated with *Bdnf IV* sgRNA but not with *Bdnf I* sgRNA (*n* = 8, one-way ANOVA, F_(3, 28)_ = 34.16, *P* < 0.0001). (**d**) Total *Bdnf IX* transcript levels are upregulated with both *Bdnf I* and *Bdnf IV* sgRNAs (*n* = 8, one-way ANOVA, F_(3, 28)_ = 277.7, *P* < 0.0001). sgRNA designed for the bacterial *LacZ* gene is used as a non-targeting control in panels (**b-d**). Dunnett’s *post hoc* test was used for individual comparisons. All data are expressed as mean ± s.e.m. Individual comparisons, ^**^*P* < 0.01, ^***^*P* < 0.001, ^****^*P* < 0.0001.

We designed sgRNAs to target two promoter regions upstream of either *Bdnf I* or *Bdnf IV* exons. These two *Bdnf* transcripts are known to be epigenetically regulated, are responsive to neuronal stimulation, and regulate LTP and memory formation^30, 33–35^. CRISPRa targeting in hippocampal cultures to *Bdnf I* selectively increased the expression of *Bdnf I* transcript variant, which reflected in the increase of the total *Bdnf* mRNA as measured by exon *IX* upregulation (**Figure 3b,d**). Likewise, co-transduction of dCas9-VPR and *Bdnf IV* sgRNA specifically upregulated the expression of *Bdnf IV* variant and also increased total *Bdnf IX* mRNA levels (**Figure 3c-d**). Multiplexing both sgRNAs for *Bdnf I* and *IV* drove the expression of both transcript variants and produced a maximal upregulation of total *Bdnf IX* levels (**Figure 3b-d**). Using *Bdnf* transcript variant manipulation, our data demonstrate specificity of the CRISPRa system at an individual mRNA transcript level.

### Transcriptome-wide selectivity of CRISPRa

CRISPR-based targeting relies on complementary sequence identity between the sgRNA and genomic DNA. Therefore, off-target sgRNA binding and gene induction is possible if there is sufficient sequence similarity^36^. To evaluate specificity with *Bdnf* transcript induction, we performed whole-transcriptome RNA-seq after CRISPRa targeting of *Bdnf I* or *IV* in hippocampal cell cultures. Quantification of transcript abundance (using fragments per kilobase per million mapped reads (FPKM) values) for each non-coding *Bdnf* exon(*I* – *VIII*) and the common-coding exon *IX* revealed that targeting either exon *I* or *IV* increased the respective transcript variant without altering adjacent transcripts. Targeting either exon *I* or *IV* also increased the abundance of the coding *Bdnf IX* exon (**Figure 4a-b**). Although *Bdnf I* or *Bdnf IV* sgRNA sequences were completely unique within the rat genome assembly (with no complete matches elsewhere), it was possible that CRISPRa could induce off-target effects at other genes. To examine this, we performed an extensive algorithmic search for potential off-target DNA sequences using Cas-OFFinder^37^, allowing systematic identification of similar sequences with up to 4 nucleotide mismatches to our sgRNAs (see **Supplementary Tables 2** and **3** for complete list). Most potential off-target loci fell within intergenic regions distant from any annotated genes. However, even for predicted off-target sites located within or near genes (+/- 2 kbp), we detected few gene expression changes with either sgRNA manipulation. For *Bdnf I* CRISPRa targeting, we identified 61 predicted off-target genes (annotated in orange in **Figure 4c**), but only 7 (11.5%) were significantly altered as compared to the *LacZ* control group (4 upregulated genes and 3 downregulated genes). Likewise, for *Bdnf IV* sgRNA targeting, we identified 23 predicted off-target genes (**Figure 4d**), only 6 (26.1%) of which were differentially expressed genes (3 upregulated genes and 3 downregulated genes versus *LacZ* controls). Given that the percentages of predicted off-target genes significantly altered in each case were similar to the overall percentage of genes altered in *Bdnf I* and *Bdnf IV* CRISPRa targeting (5.3% and 22.9%, respectively), and that observed changes included both increases and decreases in gene expression, we interpret these results to indicate a lack of direct off-target effects using CRISPRa. Finally, genes directly upstream and downstream of *Bdnf* on the third chromosome (*Lin7c* and *Kif18a*) were not differentially expressed following either manipulation, suggesting that on-target effects do not alter the expression of nearby genes. Together, these results illustrate the selectivity of the CRISPRa system, which robustly upregulated the expression of select transcript variants of *Bdnf* without driving adjacent genes or predicted off-target loci.

**Figure 4.**
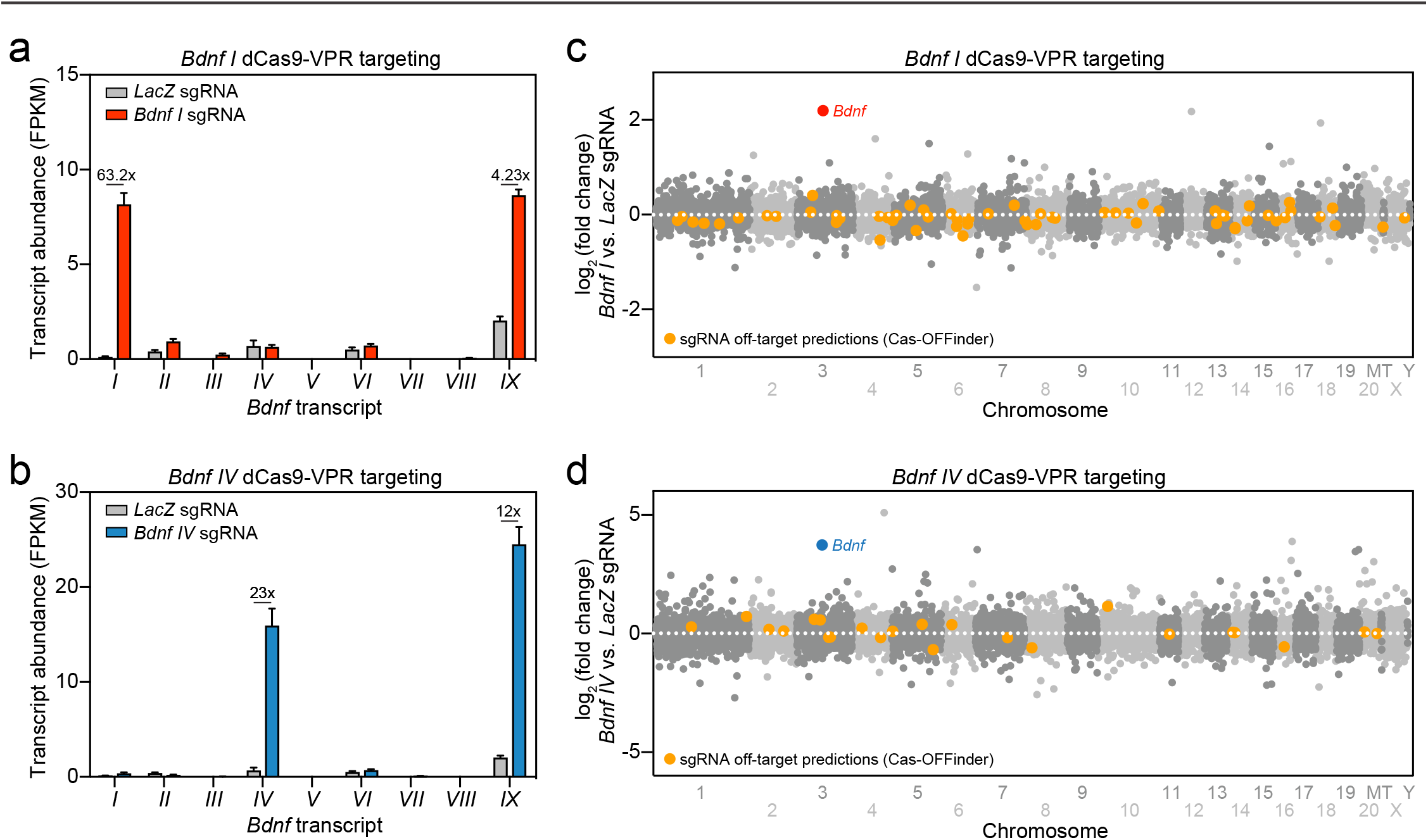
Transcriptome-wide selectivity of CRISPRa at *Bdnf* non-coding exons and the absence of off-target gene upregulation revealed by RNA-seq. (**a-b**) *Bdnf* transcript variant expression (FPKM values) following dCas9-VPR targeting with *Bdnf I* (**a**) and *Bdnf IV* (**b**) sgRNAs. *Bdnf I* sgRNA treatment upregulated *Bdnf I* transcripts by 63.2x (**a**), while *Bdnf IV* sgRNA treatment upregulated *Bdnf IV* transcripts by 23x (**b**). Both *Bdnf I* and *IV* sgRNA targeted conditions increased *Bdnf IX* transcript expression by 4.23x and 12x, respectively. sgRNA designed for the bacterial *LacZ* gene is used as a non-targeting control. (**c-d**) Mirrored Manhattan plots showing degree of mRNA change across the genome for *Bdnf I* (**c**) and *Bdnf IV* (**d**) dCas9-VPR targeting. While there were no exact matches for *Bdnf I* or *Bdnf IV* sgRNA sequences elsewhere in the genome, all potential off-target sites with up to 4 nucelotide mismatches (identified with Cas-OFFinder) are shown in orange.

### Downstream transcriptional outcomes following CRISPRa at Bdnf

To investigate the identity of genes differentially regulated by *Bdnf I* or *IV* upregulation using CRISPRa, we first characterized differentially expressed genes (DEGs) in either *Bdnf I* or *IV* versus *LacZ* targeted conditions. In both datasets, *Bdnf* was the top significantly upregulated gene (**Figure 5a-b**). We detected 387 upregulated genes and 277 downregulated genes after *Bdnf I* induction as well as 1651 upregulated genes and 1191 downregulated genes after *Bdnf IV* targeting (**Figure 5c-d**). Out of the 664 DEGs altered by *Bdnf I* upregulation and 2842 DEGs altered by *Bdnf IV* upregulation, 259 genes were shared in both conditions (**Figure 5e**). At these 259 co-regulated genes, nearly all (238 of 259, 91.9%) were regulated in the same direction by *Bdnf I* and *Bdnf IV* targeting. Increased *Bdnf* levels were associated with elevated expression of several IEGs that are often used as markers for neuronal activation, including *Arc*, *Fos*, *Egr1*, and *Egr3* (**Figure 5f**). These results complement previous studies linking *Bdnf* signaling with IEG expression^38,39^.

**Figure 5.**
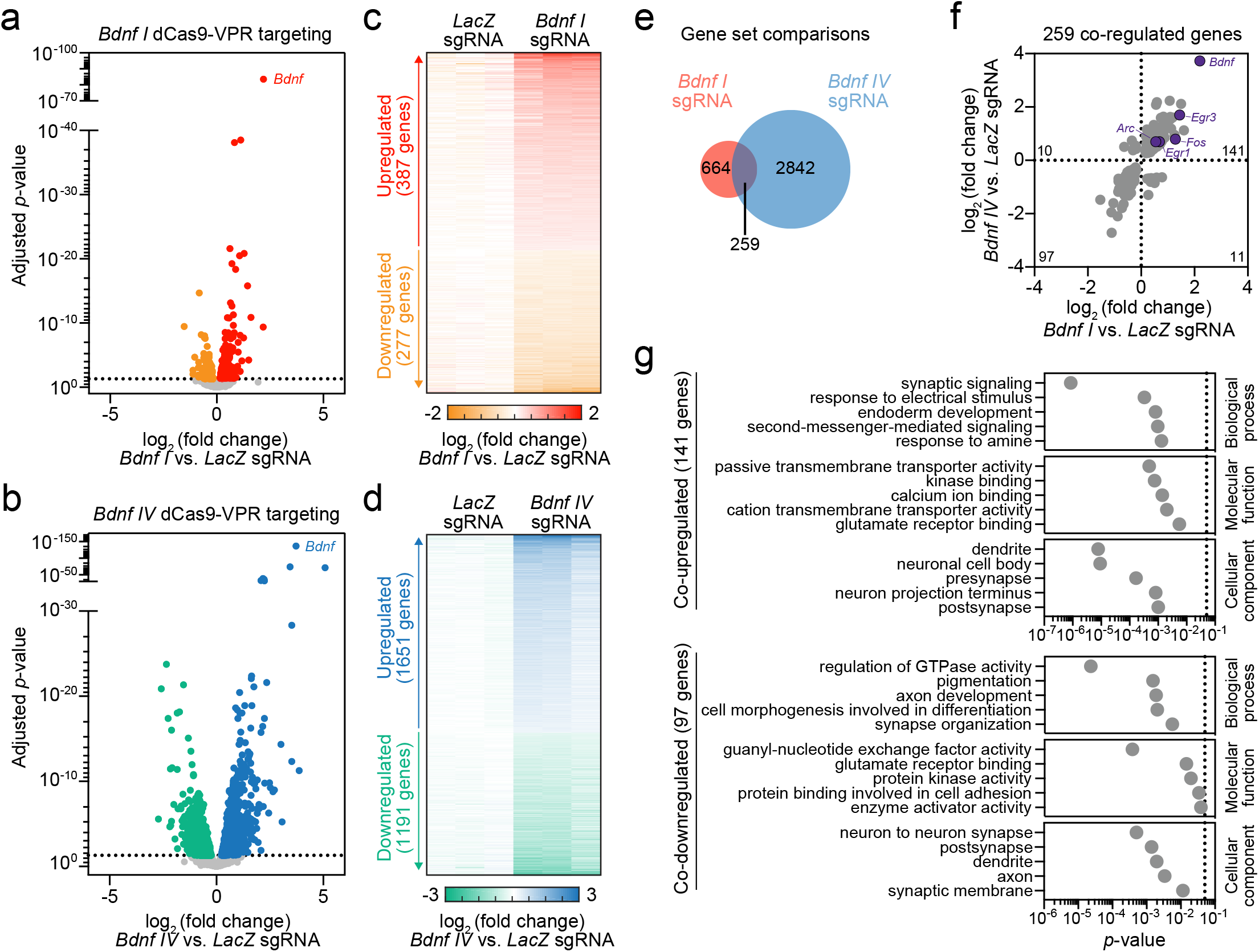
CRISPRa targeted induction of *Bdnf I* and *IV* transcript variants causes coordinated upregulation of genes involved in neuronal activation and synaptic function. (**a-b**) RNA-seq volcano plots showing differentially expressed genes (DEGs) detected by DESeq2 in *LacZ* vs. *Bdnf I* sgRNA (**a**) and *LacZ* vs. *Bdnf IV* sgRNA (**b**) targeted conditions. Standard cutoff point is represented by the horizontal dotted line (adjusted *P* < 0.05). Upregulated (red or blue) and downregulated (orange or green) genes are indicated for each comparison. *Bdnf* is the top upregulated gene in both conditions. (**c-d**) Heat maps representing all DEGs comparing *LacZ* vs. *Bdnf I* sgRNA (**c**) and *LacZ* vs. *Bdnf IV* sgRNA (**d**) targeted conditions for three biological replicates. Values in each row represent *LacZ*-normalized counts for each DEG (adjusted *P* < 0.05). Log_2_ fold change increases (red or blue) or decreases (orange or green) in gene expression are presented relative to the *LacZ* mean (white). (**e**) Venn diagram representing 664 DEGs after *Bdnf I* sgRNA targeting (red) and 2,842 DEGs after *Bdnf IV* sgRNA targeting (blue), with 259 overlapping genes. (**f**) Scatter plot representing all shared 259 DEGs in *Bdnf I* vs. *Bdnf IV* sgRNA targeted conditions. Genes upregulated in both groups (141), downregulated in both groups (97), upregulated after *Bdnf I* and downregulated after *Bdnf IV* sgRNA targeting (11), downregulated after *Bdnf I* and upregulated after *Bdnf IV* sgRNA targeting (10) are indicated. Select upregulated IEGs are specified. (**g**) Top significant gene ontology (GO) terms for 141 co-upregulated and 97 co-downregulated genes in *Bdnf I* and *Bdnf IV* sgRNA targeted conditions.

Gene ontology (GO) analysis revealed co-upregulated genes shared by both *Bdnf I* and *IV*-targeting conditions were enriched for synaptic signaling, response to stimulation, and second-messenger signaling activation (**Figure 5g**, top panel). Additionally, co-upregulated genes are enriched in molecular functions ranging from transmembrane transporter activity to kinase and glutamate receptor binding and are enriched for synaptic and projection-specific compartmentalization (**Figure 5g**, top panel). Genes that were co-downregulated are involved in the regulation of signaling molecule activity, cell differentiation, and axonal development processes (**Figure 5g**, bottom panel). Overall, the transcriptome-wide characterization of *Bdnf*-induced DEGs supports the role of *Bdnf* function in synaptic plasticity, neuronal signaling, response to glutamate, and activation of second-messenger systems^33,39^. This further highlights how CRISPRa can be used to drive gene expression profile changes to explore downstream molecular consequences of altered neuronal signaling.

### Physiological alterations following CRISPRa-mediated Bdnf and Reln upregulation

It is well established that *Bdnf* signaling enhances synaptic communication and facilitates the induction of LTP ^33,39^, and our RNA-seq results revealed that *Bdnf* induced by CRISPRa increases expression of genes commonly linked to neuronal activation. Therefore, we tested whether *Bdnf* upregulation using CRISPRa influences physiological properties of neuronal cultures. Primary hippocampal neurons were seeded directly on multi-electrode arrays (MEAs) in cell culture plates and transduced with lentiviruses expressing sgRNAs (*LacZ* control or *Bdnf I* and *IV*) and CRISPRa machinery (**Figure 6a**). For these experiments, we chose to pool *Bdnf I* and *IV* sgRNAs since that manipulation resulted in the most robust increase in the total *Bdnf IX* levels (**Figure 3b-d**). Following neuronal transduction on DIV 4, we verified expression of sgRNA lentiviral vectors using mCherry expression and performed electrophysiological recordings on DIV 7, 9, and 11 (**Figure 6 a-b**). Compared to the non-targeting control (*LacZ* sgRNA), treatment with *Bdnf I* and *IV* sgRNAs increased action potential frequency in the top one-third most active neurons by DIV 11 without changing the number of active units across the two conditions (**Figure 6 c-f**). In addition, the frequency of action potential bursts was increased, indicating selective communication between neurons and a greater potential for enhanced synaptic plasticity (**Figure 6g**). Following electrophysiological recordings, we verified efficient CRISPRa at *Bdnf IX* using RT-qPCR on RNA extracted from individual culture wells (**Figure 6h**). Collectively, these experiments demonstrate that upregulation of *Bdnf* gene expression using CRISPRa increases baseline neuronal activity patterns.

**Figure 6.**
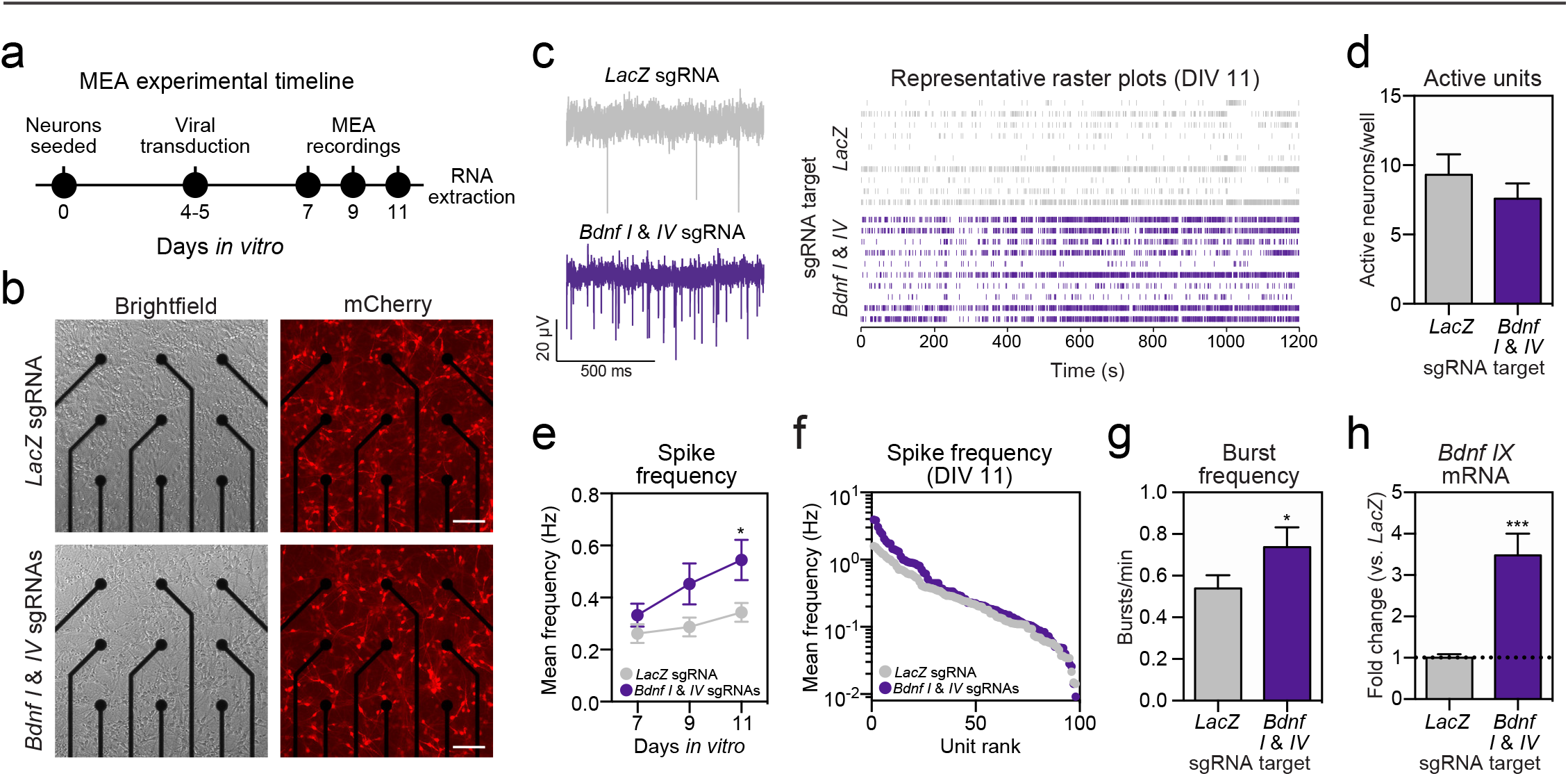
CRISPRa induction of *Bdnf* mRNA increases spike and burst frequency in hippocampal neurons cultured on microelectrode arrays (MEAs). (**a**) Experimetnal timeline for viral transduction, MEA recordings, and RNA extraction. (**b**) Primary hippocampal neurons grown on MEAs and transduced with dCas9-VPR and *LacZ* (top) or *Bdnf I* & *IV* (bottom) sgRNAs. mCherry signal indicates successful transduction of sgRNAs in live cultures (right). Scale bar = 100 μm. (**c**) Representative traces (left) and raster plots from 10 units (right) after *LacZ* (top) or *Bdnf I* and *IV* (bottom) targeting. (**d**) The number of active units per well does not change between *LacZ* and *Bdnf I* and *IV* - targeted conditions (*n* = 10 - 12, unpaired Student’s *t*-test; *P* = 0.1783). (**e**) Action potential frequency across DIV 7 - 11 showing an increase of mean frequency after *Bdnf I* and *IV* sgRNA treatment by DIV 11, as compared to *LacZ* sgRNA (*n* = 57 - 98 neurons, two-way ANOVA with main effect of sgRNA, F_(1, 493)_ = 8.561, *P* = 0.0036, Sidak’s *post hoc* test for multiple comparison). (**f**) Spike frequency at DIV 11 for all units ranked from highest to lowest mean frequency showing an increase in activity for the top 1/3 most active units in *Bdnf I* and *IV* vs. *LacZ* targeted conditions. (**g**) Burst frequency at DIV 11 is increased after *Bdnf I* and *IV* vs. *LacZ* targeting (*n* = 98, unpaired Student’s *t*-test; *P* = 0.0392). (**h**) Upregulated *Bdnf IX* mRNA expression after *Bdnf I* and *IV* vs. *LacZ* targeting following MEA recordings (*n* = 10 - 12, unpaired Student’s *t*-test; *P* = 0.0002). All data are expressed as mean ± s.e.m. ^*^*P* < 0.05 and ^***^*P* < 0.001.

To extend these observations to a second gene, we investigated neuronal activity patterns after CRISPRa-mediated upregulation of the *Reln* gene, which codes for Reelin, a large and multifunctional extracellular protein. Bidirectional modulation of *Reln* expression has been shown to affect neuronal function and synaptic activity by altering the NDMA receptor^40,41^. Additionally, the *Reln* locus is large, taking up approximately 426 kbp of genomic DNA, making it a difficult target for traditional genetic manipulations such as cDNA overexpression cassettes. In cultured hippocampal neurons plated on MEAs and recorded on DIV 7, we found that wells containing the *Reln*-targeted dCas9-VPR construct were not functionally distinct from controls in that there was not a significant difference in action potential frequency or bursting activity (**Supplementary Figure 2a-c**). However, unlike *Bdnf* manipulation, upregulation of *Reln* increased the number of spontaneously active neurons. Overall, these findings suggest a possible increase in neuronal maturation as a result of increased *Reln* expression, resulting in more physiologically active neurons.

### CRISPRa gene targeting results in increased protein levels in vivo

To examine the efficiency of the CRISPRa system in vivo, we stereotaxically infused CRISPRa lentivirus and sgRNA lentiviruses (non-targeting *LacZ* control or rat *Fosb*) into opposite hemispheres of the dorsal hippocampus, nucleus accumbens, or prefrontal cortex of adult rats (**Figure 7a-c**). After two weeks to allow for viral expression, animals were perfused and immunohistochemistry (IHC) was performed for Fosb to determine if CRISPRa targeting results in increases in protein levels. Since the mCherry signal survives fixation and does not need to be amplified with an antibody in IHC, we were able to observe the viral spread in all targeted brain regions, noting that there was robust expression of the sgRNA construct in each region regardless of *LacZ* or *Fosb* targeting. Importantly, Fosb protein expression was strongly increased in hemispheres only receiving *Fosb* sgR-NAs paired with dCas9-VPR (**Figure 7a-c**, *LacZ* targeting left, *Fosb* targeting right), indicating that increases in gene expression directly result in an increased number in Fosb+ cells in all regions (**Figure 7d-f**). These results offer evidence that CRISPRa can be used successfully in vivo in multiple neuronal populations to achieve increases in protein translation with a single viral infusion of pooled dCas9-VPR and sgRNA lentiviruses in the adult brain.

**Figure 7.**
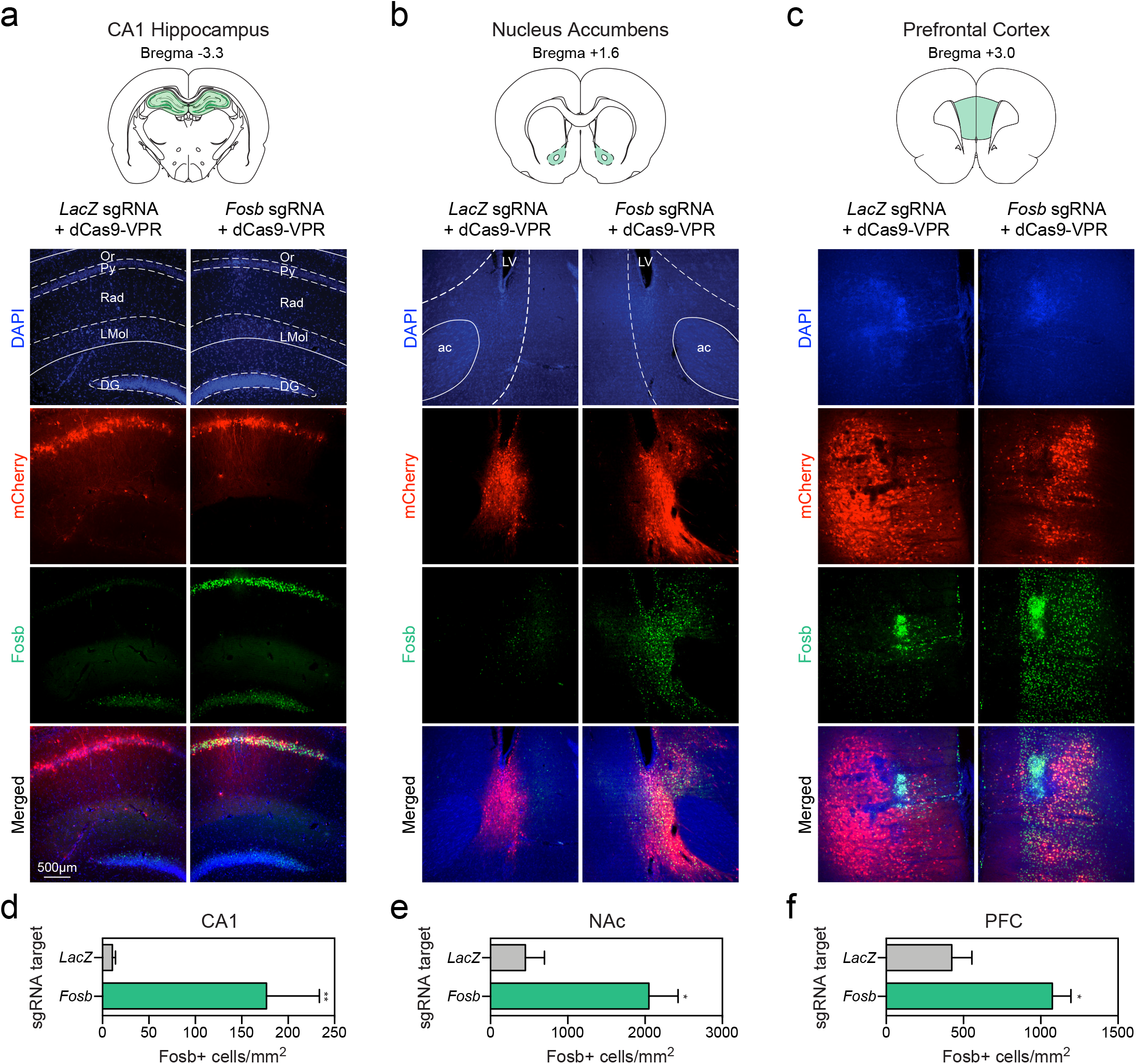
CRISPRa-mediated induction of Fosb in hippocampal, striatal, and cortical neurons *in vivo*. (**a-c**) Lentiviral infusions were bilaterally targeted to the brain region of interest^56^ in adult male rats (*n* = 4 rats/region). Two weeks following stereotaxic viral infusions, animals were transcardially perfused and immunohistochemistry (IHC) was performed to measure *Fosb* upregulation. IHC reveals high transduction efficiency of the guide RNA (expressing mCherry, signal not amplified) bilaterally in (**a**) the CA1 region of the dorsal hippocampus, (**b**) the nucleus accumbens core (NAc), and (**c**) the medial prefrontal cortex (PFC). Fosb protein is enhanced in the hemisphere that was infused with the *Fosb*-targeting sgRNA (right) compared to the hemisphere that received a sgRNA targeting the bacterial *LacZ* gene (left). Cell nuclei were stained with DAPI. Scale bar, 500 μm. Schematics of target regions are adapted from Paxinos and Watson. (**d-f**) dCas9-VPR increases the number of Fosb+ cells in the CA1, NAc, and PFC, compared to a non-targeting control (*LacZ*). (*n* = 4, ratio paired t-test; CA1: *t*_3_ = 8.73, *P* = 0.003, *R*^2^ *=* 0.96; NAc: *t*_3_ = 4.62, *P* = 0.019, *R*^2^ = 0.87; PFC: *t*_3_ = 3.43, *P* = 0.041, *R*^2^ = 0.79). All data are expressed as mean ± s.e.m. Individual comparisons, ^*^*P* < 0.05 and ^**^*P* < 0.01. Or: oriens layer, Py: pyramidal cell layer, Rad: radiatum layer, LMol: lacunosum moleculare, DG: dentate gyrus, ac: anterior commissure, LV: lateral ventricle.

## DISCUSSION

Unraveling transcriptional control of specific neuronal properties and functions requires tools that can achieve robust, selective, and modular induction of gene expression. Here, we present a neuron-optimized CRISPRa system capable of inducing targeted gene expression in post-mitotic neurons. This system allows efficient targeting of a wide variety of genes that are critical for neuronal processes, including genes of various lengths, cellular roles, and physiological functions. We demonstrate that this optimized CRISPRa system is effective in multiple neuronal populations, including cortical, hippocampal, and striatal neurons both *in vitro* and *in vivo*. Moreover, multiplexed pooling of sgRNAs enables synergistic upregulation of a single target or coordinated control over many genes. We highlight the unprecedented selectivity of the CRISPRa system by driving the expression of individual *Bdnf* mRNA transcript variants without globally affecting non-targeted variants or off-target genes, as well as the utility of this system for studying how single-gene manipulations alter gene expression programs and neuronal physiology. Together, these results provide compelling support for application of CRISPRa approaches to the study of gene regulation in diverse neuronal systems.

A key limitation to current gene overexpression approaches is the inability to express long genes using common viral vectors such as AAVs or lentiviruses. Our neuron-optimized lentivirus-based CRISPRa system provides an opportunity to expand the number of possible genetic screens in the CNS, especially for genes that are too long to be packaged in an overexpression vector. In this study, we successfully targeted genes of variable lengths: shorter genes such as *Ascl1* (1.8 kbp) and *Fos* (2.8 kbp), medium-length genes such as *Bdnf* (50 kbp), and longer genes such as *Reln* (426 kbp) and *Ebf1* (389 kbp). While typical overexpression systems would require increased viral capacity to express long genes, this CRISPRa system has a fixed cargo size given that sgRNA length does not need to increase with gene size. Importantly, this lentiviral-mediated construct delivery system allows for transgene expression within one week *in vitro* and two weeks *in vivo* (**Figures 1 & 7**), while also providing stable genome integration for potentially long-lasting upregulation. Additionally, the greater packaging capacity of the lentiviral capsid (~10 kbp) is ideal for the larger dCas9-VPR construct, as opposed to other viral vectors with lower packaging capacity, such as an AAV (~4.7 kbp)^42^. Moreover, these lentivirus-compatible constructs can be packaged into high-titer lentiviruses capable of high neuronal efficiency. Thus, this system can be used to drive a variety of genes regardless of length or complexity in post-mitotic neurons.

While the emergence of next-generation sequencing has allowed for unprecedented insight into the genome-wide changes in gene expression during development or in response to environmental stimuli, methods to mimic larger-scale gene expression profiles have been lacking. With CRISPRa, simultaneous activation of multiple gene targets allows for the investigation of global transcriptomic states, in addition to candidate gene approaches. At the *Fos* and *Fosb* genes, we found that pooling multiple sgRNAs drove more robust increases in gene expression, potentially enabling gene expression changes to be carefully and stably titrated to achieve alterations that mimic physiological conditions. Likewise, we found that multiplexing sgRNAs across genes enabled simultaneous expression of genes that are often co-regulated by neuronal depolarization, enabling more effective experimental dissection of cooperative gene programs that link neuronal activation to long-term adaptive changes.

Despite using the same dCas9-VPR fusion as a transcriptional activator at all genes, we found remarkable variability in levels of gene induction following CRISPRa. This variability is likely influenced by multiple factors, including sgRNA placement relative to gene regulatory elements, chromatin accessibility, and baseline gene expression levels^18,19,43^. In combination with rapidly growing transcriptome- and genome-wide datasets from distinct neuronal structures and subtypes, it is likely that these factors can be effectively harnessed to establish predictable rules for gene induction across neuronal systems. Similarly, we anticipate that this approach can easily be expanded to incorporate other fusion proteins, such as gene repressors or enzymes that catalyze or remove histone and DNA modifications. Indeed, using a previously neuron-optimized CRISPRi system, we also found that some sgRNAs can be repurposed for bidirectional modulation of gene expression, demonstrating the flexibility and modular nature of this approach.

The CRISPRa system allows for the investigation of unique biological questions not feasible to study using other approaches. For example, the functional significance of exon-specific promoter usage during transcription of *Bdnf* has been a long-standing question in the field of neuroscience^25^. Differential expression of diverse *Bdnf* transcript variants have been described in numerous physiological states, such as development and adult synaptic plasticity, as well as neurodevelopmental and psychiatric disorders such as addiction, schizophrenia, and depression^44^. Here, we demonstrate exquisite selectivity of CRISPRa at a single transcript variant of *Bdnf* while leaving non-targeted *Bdnf* transcripts and potential off-target genes unaffected. RNA-seq analysis after specific *Bdnf* variant upregulation showed an enhancement of genes involved in synaptic plasticity, neuronal excitability and dendritic arborization, all consistent with the known roles of *Bdnf* in the nervous system^33^. Upregulation of *Bdnf* gene expression lead to an increase in spike and burst frequency in cultured hippocampal neurons, further supporting previous reports that Bdnf can potentiate synaptic plasticity^25,33,45^. This example illustrates how CRISPRa could be used to investigate the function of not only individual genes, but also diverse transcript variants of genes in complex neuronal systems.

An additional advantage of this CRISPRa approach is the ease of transfer across model systems. In our studies, we utilized the outbred Sprague Dawley rat strain. While this organism is commonly used to model complex behavioral and cognitive processes and is often viewed to have more relevance as a model of human disease^46^, it has not been as readily amenable to genetic manipulations as *D. melanogaster*, *C. elegans,* or mouse model systems. This drawback has led to generation of fewer transgenic rat lines, which delays incorporation of this important model system into investigations targeting molecular mechanisms. This newly-optimized CRISPRa system provides more avenues for mechanistic work in rats and other model species.

This CRISPRa system is comprised of a constitutively active construct. Adaptation of these CRISPRa tools to an inducible system or viral systems with more transient expression will allow further flexibility of use and precise temporal control of gene expression. For example, during development, temporal gene expression is critical to establish cell type and proper connectivity in the developing brain. In adulthood, neuronal activity alters cellular signaling cascades, which often converge in the nucleus to alter gene expression as a result of environmental stimulation. To gain even tighter temporal control on transcription, this system could be adapted into existing chemical or physical inducible systems^20^. Additionally, while this study did target specific neuronal subpopulations with subpopulation-associated promoters (excitatory, inhibitory, and modulatory neuron-associated promoters), this addition could enable powerful circuit-specific targeting through use of cell-type specific promoters or Cre transgenic animals.

In short, here we establish a robust and neuron-optimized CRISPR/dCas9 activator system for specific upregulation of gene expression. The CRISPRa system is fast, inexpensive, modular, and drives potent and titratable gene expression changes from the endogenous gene loci in vivo and in vitro, making it more advantageous over traditional genetic manipulations, such as the use of transgenic animals or overexpression vectors. We propose that the CRISPRa system will be a readily accessible tool for the use in the investigation of gene function in the central nervous system.

## METHODS

### Animals

All experiments were performed in accordance with the University of Alabama at Birmingham Institutional Animal Care and Use Committee. Sprague-Dawley timed pregnant dams and 90-120-day-old male rats were purchased from Charles River Laboratories. Dams were individually housed until embryonic day 18 (E18) for cell culture harvest, while male rats were co-housed in pairs in plastic cages in an AAALAC-approved animal care facility on a 12-hour light/ dark cycle with *ad libitum* food and water. Animals were randomly assigned to experimental groups.

### Neuronal Cell Cultures

Primary rat neuronal cultures were generated from E18 rat cortical, hippocampal, or striatal tissue as described previously^47,48^. Briefly, cell culture plates (Denville Scientific Inc.) and MEAs (Multichannel Systems) were coated overnight with poly-L-lysine (Sigma-Aldrich; 50 μg/ml) and rinsed with diH_2_O. Hippocampal and striatal culture plates were supplemented with 7.5 μg/mL laminin (Sigma-Aldrich). Dissected cortical, hippocampal, or striatal tissue was incubated with papain (Worthington LK003178) for 25 min at 37°C. After rinsing in complete Neurobasal media (supplemented with B27 and L-glutamine, Invitrogen), a single cell suspension was prepared by sequential trituration through large to small fire-polished Pasteur pipettes and filtered through a 100 μm cell strainer (Fisher Scientific). Cells were pelleted, re-suspended in fresh media, counted, and seeded to a density of 125,000 cells per well on 24-well culture plates (65,000 cells/cm^2^) or 6-well MEA plates (325,000 cells/cm^2^). Cells were grown in complete Neurobasal media for 11 days in vitro (DIV 11) in a humidified CO_2_ (5%) incubator at 37°C with half media changes at DIV 1, 4-5, and 8-9. MEAs received a one-half media change to BrainPhys (Stemcell Technologies Inc.) with SM1 and L-glutamine supplements starting on DIV 4-5 and continued every 3-4 days.

### RNA extraction and RT-qPCR

Total RNA was extracted (RNAeasy kit, Qiagen) and reverse-transcribed (iScript cDNA Synthesis Kit, Bio-Rad). cDNA was subject to RT-qPCR for genes of interest, as described previously^48^. A list of PCR primer sequences is provided in **Supplementary Data Table 1**.

### CRISPR-dCas9 construct design

For transcriptional activation, a lentivirus-compatible backbone (a gift from Feng Zhang, Addgene #52961)^49^ was modified by insertion of dCas9-VPR (VP64-p65-Rta) cassette driven by one of the following promoters: EF1α (human elongation factor 1 alpha), PGK (human phosphoglycerate kinase), CAG, and SYN (human synapsin 1 promoter). SP-dCas9-VPR was a gift from George Church (Addgene #63798)^28^. For transcriptional repression, the SYN promoter was cloned into the lentivirus compatible KRAB-dCas9 construct, which was a gift from Jun Yao^22^. A guide scaffold (a gift from Charles Gersbach, Addgene #47108)^50^ was inserted into a lentivirus compatible backbone, and EF1a-mCherry was inserted for live-cell visualization. A *BsmBI* cut site within the mCherry construct was mutated with a site-directed mutagenesis kit (NEB). Gene-specific sgRNA targets were designed using online tools provided by the Zhang Lab at MIT (crispr.mit.edu) and CHOPCHOP (http://chopchop.cbu.uib.no/). To ensure specificity all CRISPR RNAs (crRNAs), sequences were analyzed with National Center for Biotechnology Information’s (NCBI) Basic Local Alignment Search Tool (BLAST). A list of the target sequences is provided in **Supplementary Table 1**. crRNAs were annealed and ligated into the sgRNA scaffold using the *BsmBI* cut site. Plasmids were sequence-verified with Sanger sequencing. The bacterial *LacZ* gene target was used as a sgRNA non-targeting control.

### Transfection

HEK293T cells were obtained from American Type Culture Collection (ATCC CRL-3216) and were maintained in DMEM + 10% FBS. Cells were seeded at 80k in 24 well plates the day before transfection, and 500ng of plasmid DNA was transfected in molar ratio (sgRNA:dCas9-VPR) with FuGene HD (Promega) for 40 hrs before RNA extraction and downstream RT-qPCR analysis.

### Nucleofection

C6 cells were obtained from American Type Culture Collection (ATCC CCL-107) and cultured in F-12k-based medium (2.5% bovine serum, 12% horse serum). At each passage, cells were processed for nucleofection (2 x10^6^ cells/group). Cell pellets were resuspended in nucleofection buffer (5 mM KCl, 15 mM MgCl, 15 mM HEPES, 125 mM Na_2_HPO_4_/NaH_2_PO_4_, 25 mM mannitol) and nucleofected with 3.4 μg plasmid DNA per group. Nucleofector^™^2b device (Lonza) was used according to the manufacturer’s instruction (C6, high efficiency protocol). Nucleofection groups were diluted with 500 μl media and plated in triplicates in 24-well plates (~666,667 cells/well). Plates underwent a full media change 4-6 hrs after nucleofection, and were imaged and processed for RT-qPCR after 16 hrs.

### Lentivirus production

Large scale viruses: Viruses were produced in a sterile environment subject to BSL-2 safety by transfecting HEK-293T cells with the specified CRISPR plasmid, the psPAX2 packaging plasmid, and the pCMV-VSV-G envelope plasmid (Addgene 12260 & 8454) with FuGene HD (Promega) for 40-48 hrs in supplemented Ultraculture media (L-glutamine, sodium pyruvate, and sodium bicarbonate) in either a T75 or T225 culture flask. Supernatant was passed through a 0.45 μm filter and centrifuged at 25,000 rpm for 1 hr 45 min at 4°C. The viral pellet was resuspended in 1/100^th^ supernatant volume of sterile PBS and stored at −80°C. Physical viral titer was determined using Lenti-X qRT-PCR Titration Kit (Takara), and only viruses greater than 1×10^9^ GC/ml were used. Viruses were stored in sterile PBS at −80°C in single-use aliquots. For smaller scale virus preparation, each sgRNA plasmid was transfected in a 12-well culture plate as described above. After 40-48 hrs, lentiviruses were concentrated with Lenti-X concentrator (Takara), resuspended in sterile PBS, and used immediately or stored at −80°C in single use aliquots.

### ICC/IHC

Immunocytochemistry was performed as described previously^48^. To validate expression of the dCas9-VPR cassette, anti-FLAG primary antibody (1:5000 in PBS with 10% Thermo Blocker BSA and 1% goat serum, Thermo Fisher anti-FLAG MA1-91878) was incubated overnight at 4°C. Cells were washed three times with PBS and incubated for 1 hr at room temperature with a fluorescent secondary antibody (Alexa 488 goat anti-mouse, Invitrogen A-10667, 1:500). Cells were washed three times with PBS and mounted onto microscope coverslips with Prolong Gold anti-fade medium (Invitrogen) containing 4,6-diamidino-2-phenylindole (DAPI) stain as a marker for cell nuclei. For immunohistochemistry, adult male rats were transcardially perfused with formalin (1:10 dilution in PBS, Fisher). Brains were removed and post-fixed for 24 hrs in formalin, then sliced at 50 μm using a vibratome. Cells were permeabilized with 0.25% Triton-X in PBS, then blocked for 1 hr at room temperature with blocking buffer (1X PBS with 10% Thermo Blocker BSA and 1% goat serum). To quantify the number of Fosb+ cells, slices were incubated with an anti-Fosb primary antibody (Abcam ab11959, 1:1000 in PBS with 10% Thermo Blocker BSA and 1% goat serum) and processed as outlined above. 20x images of each infusion site were taken on a Nikon TiS inverted fluorescent microscope by first locating the center of the mCherry signal in the targeted region, and using this as a region of interest for imaging for Fosb immunoreactivity. Fosb+ cells were calculated from one projected Z stack per animal per brain region in ImageJ following background subtraction. Automated cell counts were obtained from each image using 3D object counter v2.0, with thresholds set at the same levels for both *LacZ* and *Fosb* sgRNA targeted regions within the same animal and between all animals with the same targeted region.

### Multi Electrode Array Recordings

Single neuron electrophysiological activity was recorded using a MEA2100 Lite recording system (Multi Channel Systems MCS GmbH). E18 rat primary hippocampal neurons were seeded in 6-well multielectrode arrays (MEAs) at 125,000 cells/ well (325,000 cells/cm^2^), as described above. Each MEA well contained 9 extracellular recording electrodes and a ground electrode. Neurons were transduced with CRISPRa constructs on DIV 4-5 and 20 min MEA recordings were performed at DIV 7, 9, and 11 while connected to a temperature-controlled headstage (monitored at 37°C) containing a 60-bit amplifier. Electrical activity was measured by an interface board at 30 kHz, digitized, and transmitted to an external PC for data acquisition and analysis in MC_Rack software (Multi Channel Systems). All data were filtered using dual 10 Hz (high pass) and 10,000 Hz (low-pass) Butterworth filters. Action potential thresholds were set manually for each electrode (typically > 4 standard deviations from the mean signal). Neuronal waveforms collected in MC_Rack were exported to Offline Sorter (Plexon) for sorting of distinct waveforms corresponding to multiple units on one electrode channel, and confirmation of waveform isolation using principal component analysis, inter-spike intervals, and auto- or cross-correlograms. Further analysis of burst activity and firing rate was performed in NeuroExplorer. Researchers blinded to experimental conditions performed all MEA analyses.

### RNA-Sequencing

RNA-Sequencing (RNA-Seq) was carried out at the Heflin Center for Genomic Science Genomics Core Laboratories at the University of Alabama at Birmingham. RNA was extracted, purified (RNeasy, Qiagen), and DNase-treated for three biological replicates per experimental condition. 1 μg of total RNA underwent quality control (Bioanalyzer), and was prepared for directional RNA sequencing using SureSelect Strand Specific RNA Library Prep Kit (Agilent Technologies) according to manufacturer’s recommendations. PolyA+ RNA libraries underwent sequencing (75 bp paired-end directional reads; ~22-38 M reads/sample) on an Illumina sequencing platform (NextSeq2000).

### RNA-Seq Data Analysis

Paired-end FASTQ files were uploaded to the University of Alabama at Birmingham’s High Performance Computer cluster for custom bioinformatics analysis using a pipeline built with snakemake^51^ (v5.1.4). Read quality, length, and composition were assessed using FastQC prior to trimming low quality bases (Phred < 20) and Illumina adapters (Trim_Galore! v04.5). Splice-aware alignment to the Rn6 Ensembl genome assembly (v90) was performed with STAR^52^ v2.6.0c. An average of 88.4% of reads were uniquely mapped. Binary alignment map (BAM) files were merged and indexed with Samtools (v1.6). Gene-level counts were generated using the featureCounts^53^ function in the Rsubread package (v1.26.1) in R (v3.4.1), with custom options (isGTFAnnotationFile = TRUE, useMetaFeatures = TRUE, isPairedEnd = TRUE, requireBothEndsMapped = TRUE, strandSpecific = 2, and autosort = TRUE). DESeq2^54^ (v 1.16.1) in R was used to perform count normalization and differential gene expression analysis with the application of Benjamini-Hochberg false discovery rate (FDR) for adjusted *p*-values. Differentially expressed genes (DEGs) were designated if they passed a *p* < 0.05 adjusted *p*-value cutoff and contained basemeans > 50. Manhattan plots were constructed in Prism (Graphpad). Predicted off-target sgRNA hits for *Bdnf I* and *Bdnf IV* sgRNAs were identified with Cas-OFFinder, using PAM settings for SpCas9 and the Rn6 genome assembly, tolerating up to 4 mismatches. All hits, as well as annotated features within 2 kbp of each off-target prediction, are listed in **Supplementary Tables 2 & 3**.

Gene ontology (GO) analysis was conducted with co-regulated genes (genes either up- or down-regulated by both *Bdnf I* and *Bdnf IV* sgRNA treatments, as compared to *LacZ* sgRNA control) using the WEB-based Gene Set Analaysis Toolkit (WebGestalt^55^). Overrepresentation enrichment analysis was performed using non-redundant terms in biological process, molecular function, and cellular component GO categories, using the protein-coding rat genome as a reference set. Enrichment analysis applied Benjamini-Hochberg correction for multiple comparisons and required a minimum of 5 genes per enriched GO term category.

### Stereotaxic Surgery

Naïve adult Sprague-Dawley rats were anaesthetized with 4% isoflurane and secured in a stereotaxic apparatus (Kopf Instruments). During surgical procedures, an anaesthetic plane was maintained with 1–2.5% isoflurane. Under aseptic conditions, guide holes were drilled using stereotaxic coordinates (all coordinates in respect to bregma^56^. CA1 dHPC: AP: −3.3 mm, ML: ±2.0 mm; NAc core: AP: +1.6 mm, ML: ±1.4 mm; mPFC: AP: +3.0 mm, ML: ±0.5 mm) to target either dorsal hippocampus CA1 region, nucleus accumbens core, or medial prefrontal cortex. All infusions were made using a gastight 30-gauge stainless steel injection needle (Hamilton Syringes) that extended into the infusion site (from bregma: CA1: −3.1 mm, NAc core: −7.0 mm, mPFC: −4.9 mm). Bilateral lentivirus microinfusions of (1.5 μl total volume per hemisphere) were made using a syringe pump (Harvard Apparatus) at a rate of 0.25 μl/min. Injection needles remained in place for 10 min following infusion to allow for diffusion. Rats were infused bilaterally with either 1.5 μl of total lentivirus mix comprised of 0.5 μl sgRNA and 1 μl dCas9-VPR viruses in sterile PBS. After infusions, guide holes were covered with sterile bone wax and surgical incision sites were closed with nylon sutures. Animals received buprenorphine and carprofen for pain management and topical bacitracin to prevent infection at the incision site.

### Statistical Analysis

Transcriptional differences from RT-qPCR experiments were compared with either an unpaired *t*-test or one-way ANOVA with Dunnett’s or Tukey’s *post-hoc* tests where appropriate. Fosb+ cell counts in immunohistochemistry experiments were compared with a ratio paired *t*-test. Statistical significance was designated at α = 0.05 for all analyses. Statistical and graphical analyses were performed with Prism software (GraphPad). Statistical assumptions(e.g., normality and homogeneity for parametric tests) were formally tested and examined via boxplots.

### Data Availability

All relevant data that support the findings of this study are available by request from the corresponding author (J.J.D.). All constructs will be deposited, along with maps and sequences, in Addgene.

## ACKNOWLEDGEMENTS

This work was supported by NIH grants DA039650, DA034681, and MH114990 (J.J.D.), DA042514 (K.E.S.), MH112304 (S.V.B), DA041778 (F.A.S.). L.I. is supported by the Civitan International Research Center at UAB. Additional assistance to J.J.D. was provided by the UAB Pittman Scholars Program. We would like to thank Charles A. Gersbach for his assistance in design and generation of CRISPR tools, Natalie Simpkins for assistance with immunohistochemistry, and all current and former Day Lab members for assistance and support. Sequencing experiments were carried out with generous assistance from Mike Crowley in the UAB Heflin Genomics Core.

## AUTHOR CONTRIBUTIONS

K.E.S., S.V.B., and J.J.D conceived of the experiments, performed experiments, and wrote the manuscript. M.E.Z., J.S.R., N.A.G., J.J.T., C.G.D., J.N.B., D.W., and F.A.S. assisted in construct design, experiments, statistical and graphical analysis, data interpretation, and/or manuscript construction and layout. L.I. generated bioinformatics pipelines and performed primary bioinformatics analysis. L.I. and J.J.D performed secondary bioinformatics analysis. J.J.D. supervised all work. All authors have approved the final version of the manuscript.

---

**Supplementary Figure 1.**
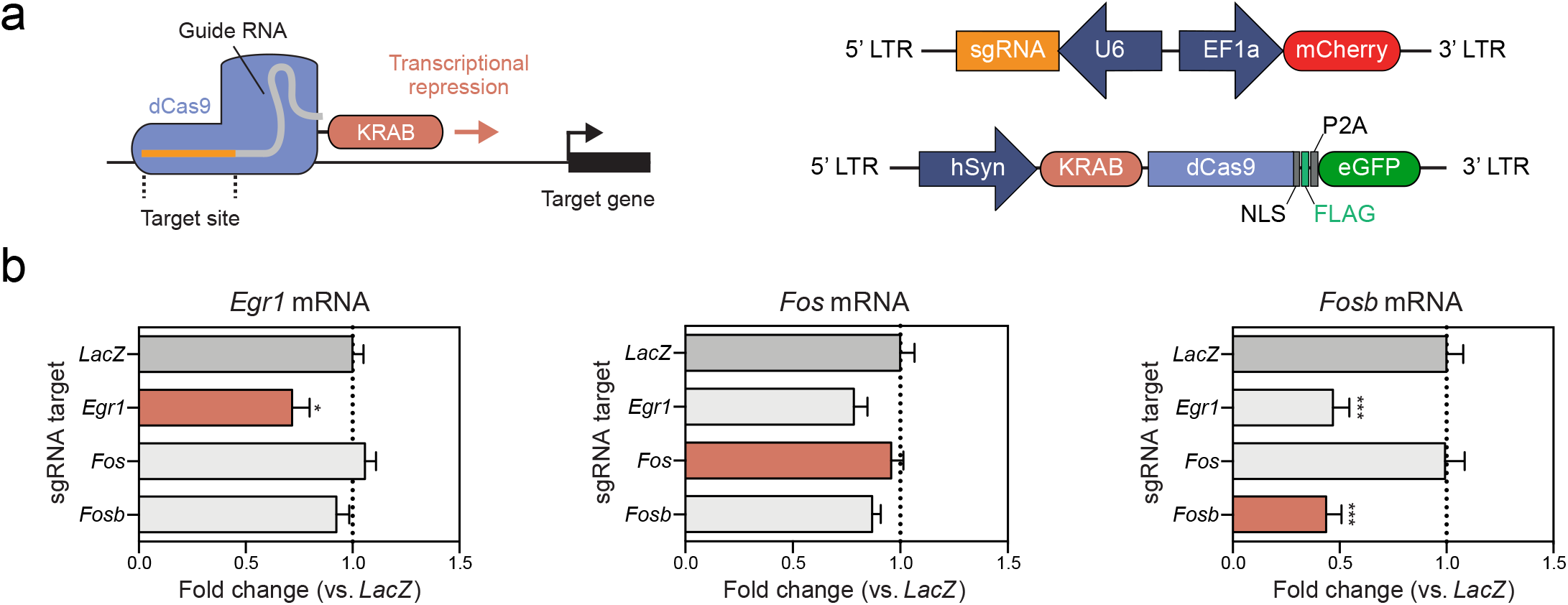
CRISPRi gene repression in primary striatal rat neurons employing the same sgRNAs utlilized with CRISPRa. (**a**) Illustration of the CRISPRi dual vector approach expressing either the single guide RNA (sgRNA) or the KRAB-dCas9. (**b**) Lentiviral transduction of primary rat striatal neurons reveals that targeting KRAB-dCas9 to the same target sites as dCas9-VPR results in gene repression of *Egr1* and *Fosb* but not *Fos* (*n* = 6, one-way ANOVA, *Egr1* F_(3,20)_ = 5.648, *P* = 0.0057; *Fos* F_(3,20)_ = 2.795, *P* = 0.0667; *Fosb* F_(3,20)_ = 15.120, *P* < 0.0001, Dunnett’s *post hoc* test for multiple comparisons). KRAB-dCas9 with a sgRNA targeted to the bacterial *LacZ* gene is used as a non-targeting control in panel (**b**). All data are expressed as mean ± s.e.m. Individual comparisons,^*^*P* < 0.05 and ^***^*P* < 0.001.

**Supplementary Figure 2.**
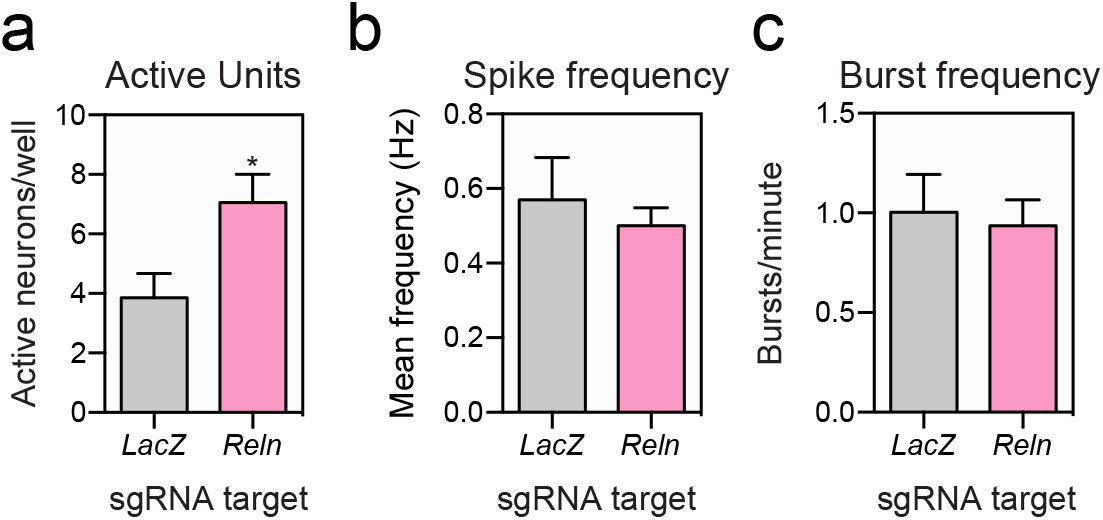
CRISPRa targeting of *Reln* in hippocampal neurons. (**a-c**) Reln targeting with CRISPRa results in more active neurons at DIV 7, but no change in spike or burst frequency (*n* = 15 wells, unpaired Student’s *t*-test; active units *t_28_* = 2.574, *P* = 0.0156). MEA recordings occured on DIV 7, approximately 72 hours after viral transduction. All data are expressed as mean ± s.e.m. Individual comparisons, ^*^*P* < 0.05.

## REFERENCES

1. Lein, E. S. et al. Genome-wide atlas of gene expression in the adult mouse brain. Nature 445, 168–176 (2007).

2. Roth, R. B. et al. Gene expression analyses reveal molecular relationships among 20 regions of the human CNS. Neurogenetics 7, 67–80 (2006).

3. Thompson, C. L. et al. A high-resolution spatiotemporal atlas of gene expression of the developing mouse brain. Neuron 83, 309–323 (2014).

4. West, A. E. & Greenberg, M. E. Neuronal activity-regulated gene transcription in synapse development and cognitive function. Cold Spring Harb Perspect Biol 3, a005744–a005744 (2011).

5. Benito, E. & Barco, A. The neuronal activity-driven transcriptome. Mol. Neurobiol. 51, 1071–1088 (2015).

6. Duke, C. G., Kennedy, A. J., Gavin, C. F., Day, J. J. & Sweatt, J. D. Experience-dependent epigenomic reorganization in the hippocampus. Learn. Mem. 24, 278–288 (2017).

7. Hermey, G. et al. Genome-wide profiling of the activity-dependent hippocampal transcriptome. PLoS ONE 8, e76903 (2013).

8. Robison, A. J. & Nestler, E. J. Transcriptional and epigenetic mechanisms of addiction. Nat Rev Neurosci 12, 623–637 (2011).

9. Jansen, R. et al. Gene expression in major depressive disorder. Mol. Psychiatry 21, 339–347 (2016).

10. Harrison, P. J. & Weinberger, D. R. Schizophrenia genes, gene expression, and neuropathology: on the matter of their convergence. Mol. Psychiatry 10, 40–68 (2005).

11. Castillo, E. et al. Comparative profiling of cortical gene expression in Alzheimer’s disease patients and mouse models demonstrates a link between amyloidosis and neuroinflammation. Scientific Reports 7, 329 (2017).

12. Prelich, G. Gene overexpression: uses, mechanisms, and interpretation. Genetics 190, 841–854 (2012).

13. Ericsson, A. C., Crim, M. J. & Franklin, C. L. A brief history of animal modeling. Mo Med 110, 201–205 (2013).

14. Fire, A. et al. Potent and specific genetic interference by double-stranded RNA in *Caenorhabditis elegans*. Nature 391, 806–811 (1998).

15. Jinek, M. et al. A Programmable Dual-RNA–Guided DNA Endonuclease in Adaptive Bacterial Immunity. Science 337, 816–821 (2012).

16. Straub, C., Granger, A. J., Saulnier, J. L. & Sabatini, B. L. CRISPR/Cas9-mediated gene knock-down in post-mitotic neurons. PLoS ONE 9, e105584 (2014).

17. Swiech, L. et al. In vivo interrogation of gene function in the mammalian brain using CRISPR-Cas9. Nature Biotechnology 33, 102–106 (2015).

18. Chavez, A. et al. Comparison of Cas9 activators in multiple species. Nat Meth 13, 563–567 (2016).

19. Konermann, S. et al. Genome-scale transcriptional activation by an engineered CRISPR-Cas9 complex. Nature 517, 583–588 (2015).

20. Savell, K. E. & Day, J. J. Applications of CRISPR/ Cas9 in the Mammalian Central Nervous System. Yale J Biol Med 90, 567–581 (2017).

21. Staahl, B. T. et al. Efficient genome editing in the mouse brain by local delivery of engineered Cas9 ribonucleoprotein complexes. Nature Biotechnology 35, 431–434 (2017).

22. Zheng, Y. et al. CRISPR interference-based specific and efficient gene inactivation in the brain. Nat Neurosci 77, 955–454 (2018).

23. Frank, C. L. et al. Regulation of chromatin accessibility and Zic binding at enhancers in the developing cerebellum. Nat Neurosci 18, 647–656 (2015).

24. Liu, X. S. et al. Editing DNA Methylation in the Mammalian Genome. Cell 167, 233–235 (2016).

25. Cunha. A simple role for BDNF in learning and memory? Frontiers in Molecular Neuroscience 3, (2010).

26. Gilbert, L. A. et al. CRISPR-Mediated Modular RNA-Guided Regulation of Transcription in Eukaryotes. Cell 154, 442–451 (2013).

27. Mali, P. et al. CAS9 transcriptional activators for target specificity screening and paired nickases for cooperative genome engineering. Nature Biotechnology 31, 833–838 (2013).

28. Chavez, A. et al. Highly efficient Cas9-mediated transcriptional programming. Nat Meth 12, 326–328 (2015).

29. Yaguchi, M. et al. Characterization of the Properties of Seven Promoters in the Motor Cortex of Rats and Monkeys After Lentiviral Vector-Mediated Gene Transfer. Human Gene Therapy Methods 24, 333–344 (2013).

30. Aid, T., Kazantseva, A., Piirsoo, M., Palm, K. & Timmusk, T. Mouse and rat BDNF gene structure and expression revisited. J. Neurosci. Res. 85, 525–535 (2007).

31. Baj, G., Leone, E., Chao, M. V. & Tongiorgi, E. Spatial segregation of BDNF transcripts enables BDNF to differentially shape distinct dendritic compartments. PNAS 108, 16813–16818 (2011).

32. An, J. J. et al. Distinct Role of Long 3’ UTR BDNF mRNA in Spine Morphology and Synaptic Plasticity in Hippocampal Neurons. Cell 134, 175–187 (2008).

33. Panja, D. & Bramham, C. R. BDNF mechanisms in late LTP formation: A synthesis and breakdown. Neuropharmacology 76, 664–676 (2014).

34. Lubin, F. D., Roth, T. L. & Sweatt, J. D. Epigenetic Regulation of Bdnf Gene Transcription in the Consolidation of Fear Memory. J Neurosci 28, 10576–10586 (2008).

35. Bredy, T. W. et al. Histone modifications around individual BDNF gene promoters in prefrontal cortex are associated with extinction of conditioned fear. Learn. Mem. 14, 268–276 (2007).

36. Sternberg, S. H., Redding, S., Jinek, M., Greene, E. C. & Doudna, J. A. DNA interrogation by the CRISPR RNA-guided endonuclease Cas9. Nature 507, 62–67 (2014).

37. Bae, S., Park, J. & Kim, J.-S. Cas-OFFinder: a fast and versatile algorithm that searches for potential off-target sites of Cas9 RNA-guided endonucleases. Bioinformatics 30, 1473–1475 (2014).

38. Cortés-Mendoza, J., Díaz de León-Guerrero, S., Pedraza-Alva, G. & Pérez-Martínez, L. Shaping synaptic plasticity: The role of activity-mediated epigenetic regulation on gene transcription. International Journal of Developmental Neuroscience 31, 359–369 (2013).

39. Bramham, C. R. & Messaoudi, E. BDNF function in adult synaptic plasticity: the synaptic consolidation hypothesis. Prog. Neurobiol. 76, 99–125 (2005).

40. Rogers, J. T. et al. Reelin supplementation enhances cognitive ability, synaptic plasticity, and dendritic spine density. Learn. Mem. 18, 558–564 (2011).

41. Chameau, P. et al. The N-terminal region of reelin regulates postnatal dendritic maturation of cortical pyramidal neurons. PNAS 106, 7227–7232 (2009).

42. Lentz, T. B., Gray, S. J. & Samulski, R. J. Viral vectors for gene delivery to the central nervous system. Neurobiol. Dis. 48, 179–188 (2012).

43. Zhou, H. et al. In vivo simultaneous transcriptional activation of multiple genes in the brain using CRISPR–dCas9-activator transgenic mice. Nat Neurosci 21, 440–446 (2018).

44. Autry, A. E. & Monteggia, L. M. Brain-derived neurotrophic factor and neuropsychiatric disorders. Pharmacol. Rev. 64, 238–258 (2012).

45. Lu, Y., Christian, K. & Lu, B. BDNF: A key regulator for protein synthesis-dependent LTP and long-term memory? Neurobiology of Learning and Memory 89, 312–323 (2008).

46. Ellenbroek, B. & Youn, J. Rodent models in neuroscience research: is it a rat race. Disease Models & Mechanisms 9, 1079–1087 (2016).

47. Day, J. J. et al. DNA methylation regulates associative reward learning. Nat Neurosci 16, 1445–1452 (2013).

48. Savell, K. E. et al. Extra-coding RNAs regulate neuronal DNA methylation dynamics. Nat Comms 7, 12091 (2016).

49. Sanjana, N. E., Shalem, O. & Zhang, F. Improved vectors and genome-wide libraries for CRISPR screening. Nat Meth 11, 783–784 (2014).

50. Perez-Pinera, P. et al. RNA-guided gene activation by CRISPR-Cas9–based transcription factors. Nat Meth 10, 973–976 (2013).

51. Köster, J. & Rahmann, S. Snakemake—a scalable bioinformatics workflow engine. Bioinformatics 28, 2520–2522 (2018).

52. Dobin, A. et al. STAR: ultrafast universal RNA-seq aligner. Bioinformatics 29, 15–21 (2012).

53. Liao, Y., Smyth, G. K. & Shi, W. featureCounts: an efficient general purpose program for assigning sequence reads to genomic features. Bioinformatics 30, 923–930 (2014).

54. Love, M. I., Huber, W. & Anders, S. Moderated estimation of fold change and dispersion for RNA-seq data with DESeq2. Genome Biol. 15, 31 (2014).

55. Wang, J., Vasaikar, S., Shi, Z., Greer, M. & Zhang, B. WebGestalt 2017: a more comprehensive, powerful, flexible and interactive gene set enrichment analysis toolkit. Nucleic Acids Research 45, W130–W137 (2017).

56. Paxinos, G. & Watson, C. The Rat Brain in Stereotaxic Coordinates. (Academic Press, 2009).

